# Fast and sensitive mapping of error-prone nanopore sequencing reads with GraphMap

**DOI:** 10.1101/020719

**Authors:** Ivan Sovic, Mile Sikic, Andreas Wilm, Shannon Nicole Fenlon, Swaine Chen, Niranjan Nagarajan

## Abstract

Exploiting the power of nanopore sequencing requires the development of new bioinformatics approaches to deal with its specific error characteristics. We present the first nanopore read mapper (GraphMap) that uses a read-funneling paradigm to robustly handle variable error rates and fast graph traversal to align long reads with speed and very high precision (>95%). Evaluation on MinION sequencing datasets against short and long-read mappers indicates that GraphMap increases mapping sensitivity by at least 15-80%. GraphMap alignments are the first to demonstrate consensus calling with <1 error in 100,000 bases, variant calling on the human genome with 76% improvement in sensitivity over the next best mapper (BWA-MEM), precise detection of structural variants from 100bp to 4kbp in length and species and strain-specific identification of pathogens using MinION reads. GraphMap is available open source under the MIT license at https://github.com/isovic/graphmap.

## Introduction

With the release of Oxford Nanopore Technologies (ONT) MinION sequencers in 2014, a new era of cheap and portable nanopore sequencers, producing ultra-long reads has become reality. Potential applications for the new technology are varied and, in addition to its use in research, its compact form factor and affordability have drawn interest for its use in point-of-care diagnostics. While some initial nanopore sequencing based applications have been reported (e.g. scaffolding and resolution of repeats in genomes^1^ and variant detection in clonal haploid samples^2^), many others remain to be explored. In particular, diploid and rare-variant calling^3^, *de novo* genome assembly^4^, metagenome assembly and pathogen identification are all promising applications that will likely see the development of new *in silico* techniques to realize them.

Read mapping and alignment tools are critical building blocks for many such applications as they help solve the difficult problem of efficiently aligning a large number of error-prone read sequences (to each other or to a reference genome) without sacrificing sensitivity or specificity. Reads from nanopore sequencing can be particularly challenging as, in addition to the volume of long reads that they generate, they also have a propensity for higher and non-uniform error profiles^5^. For example, 1D reads from the MinION sequencer have been reported to have accuracy less than 65% while a smaller fraction of high-quality (<25%; 2D) reads had accuracy greater than 70%^1^. Thus, despite the length of the reads, a sizable fraction of reads can remain unmapped (10-70%) and thus unusable for downstream applications. This is particularly the case for 1D reads which often form the bulk of the data^1,2^. While error rates continue to improve on the MinION system^2^, their variability across chemistries, sequencing runs and even within a read can be a challenge for bioinformatics pipelines. Furthermore, as new nanopore sequencing technologies become available, having a robust mapping and alignment tool that can accommodate different error profiles (i.e. ratio of insertions, deletions and substitutions) and error rates in a consistent fashion would be essential for downstream applications.

While alignment algorithms have been widely studied, gold-standard solutions such as dynamic programming (or even fast approximations such as BLAST) are too slow or infeasible in practice for aligning high-throughput sequencing reads. To address this need, a range of read mapping tools have been developed that exploit the characteristics of second-generation sequencing reads (relatively short and accurate) by trading-off a bit of sensitivity for dramatic gains in speed^6,7^. The design decisions employed in these mappers are often tuned for specific error characteristics of a sequencing technology, potentially limiting their utility across technologies and error profiles. The less than ideal results reported in recent studies using MinION data^8^ could therefore be in part due to the use of mappers (e.g. BWA-MEM^6^ and BLASR^9^) or genome aligners (e.g. LAST^10^) that are not suited to its error characteristics.

In this work, we present the first nanopore mapper (GraphMap) that is adept at mapping long and error-prone nanopore sequencing data with high sensitivity and precision. GraphMap was designed for ease-of-use, aligning reads with a wide range of lengths and error profiles without having to tune parameters. This is an important feature for a technology where error rates and profiles can vary widely across sequencing runs. Correspondingly, GraphMap also allows users to uniformly map read datasets from disparate technologies (e.g. Illumina, PacBio or ONT) with BLAST-like sensitivity and runtime comparable to state-of-the-art mappers. Experiments with several real and synthetic datasets demonstrate that GraphMap is a more sensitive mapper (than BWA-MEM, BLASR and LAST) while reporting alignments that provide highly accurate consensus sequences (Q50) with nanopore sequencing data. This in turn translates into notable advantages in real-world applications such as the use of nanopore data for single-nucleotide and structural variant calling as well as the use of MinION reads for real-time pathogen identification.

## Results

### Overview of the GraphMap algorithm

The GraphMap algorithm is structured to achieve high-sensitivity and speed using a five-stage read-funneling approach as depicted in **Figure 1a**. The underlying design principle is to have efficiently computable stages that conservatively reduce the set of candidate locations based on progressively defined forms of the read-to-reference alignment. For example, in stage I, GraphMap uses a novel adaptation of gapped spaced seeds^11^ to efficiently reduce the search space (**Figure 1b**) and get seed hits as a form of coarse alignment. These are then refined in stage II using graph-based vertex-centric processing of seeds to efficiently (allowing seed-level parallelism) construct alignment anchors (**Figure 1c**). GraphMap then chains anchors using a k-mer version of longest common subsequence (LCS) construction (stage III; **Figure 1d**), refines alignments with a form of L_1_ linear regression (stage IV; **Figure 1e**) and finally evaluates the remaining candidates to select the best location to reconstruct a final alignment (stage V). GraphMap computes a BLAST-like *E*-value as well as a mapping quality for its alignments. Further details about each of these stages can be found in the **Methods** section.

**Figure 1.**
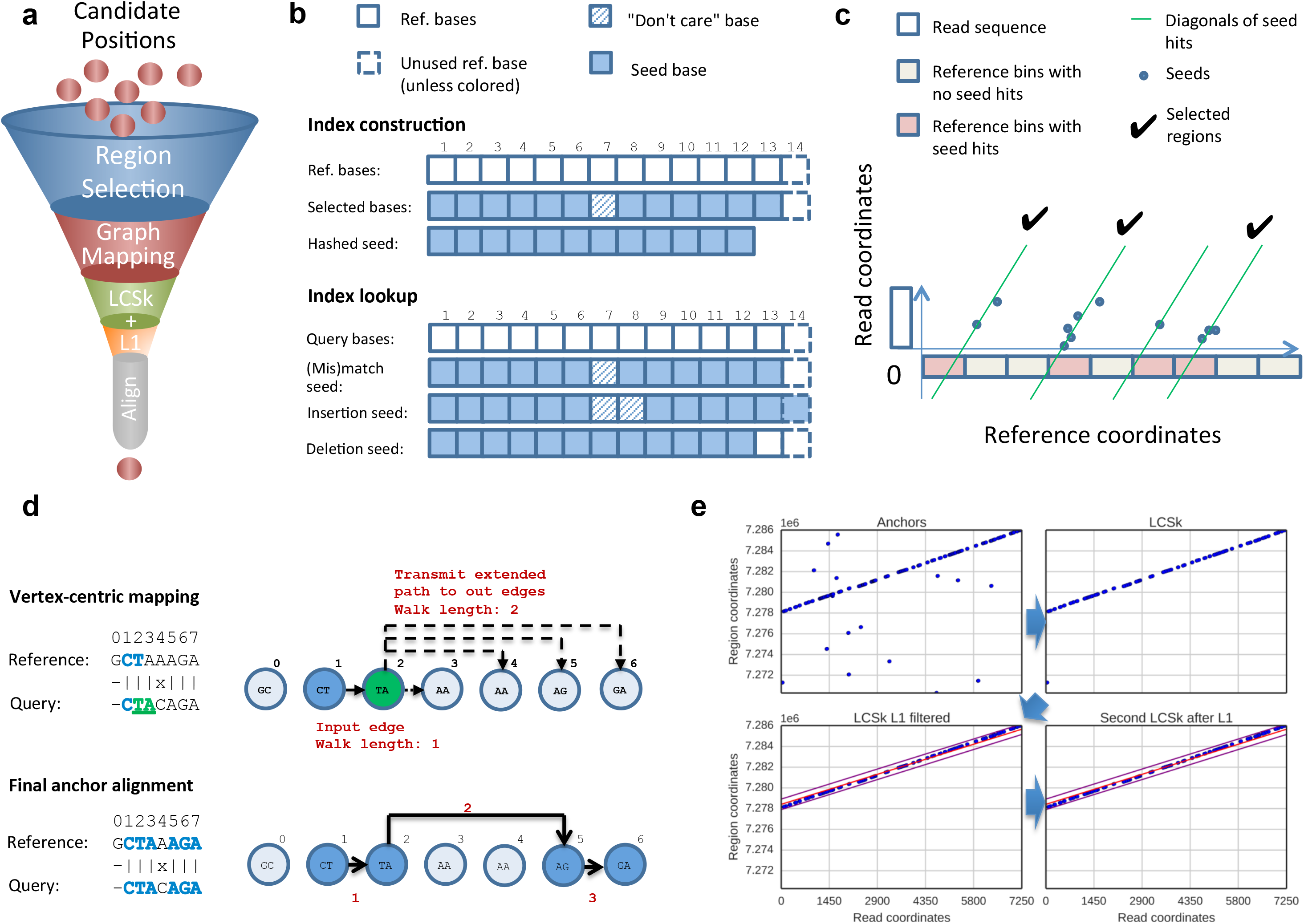
A schematic representation of stages in GraphMap. (a) Order of stages in the read-funneling approach used in GraphMap to refine alignments and reduce the number of candidate locations to one. (b) Structure of spaced seeds used for index construction and index lookup. (c) *Region selection* by clustering of candidate seeds on the reference. (d) Generation of alignment anchors through a fast graph based ordering of seeds (*Graph Mapping*). (e) Filtering of seed matches using *LCSk* search and *L1* regression.

### GraphMap maps reads accurately independent of error rates and profiles

GraphMap was designed to be efficient while being largely agnostic of error profiles and rates. To evaluate this feature, we generated a wide range of synthetic datasets that mimic the output of various sequencing technologies (Illumina, PacBio, ONT 2D, ONT 1D) and over a range of different genome sizes (**Figure 2**). We then measured GraphMaps’s precision and recall in terms of identifying the correct read *location* and in reconstructing the correct *alignment* to the reference (**Methods**). We distinguish between the two as, in principle, a mapper can identify the correct location but compute an incorrect alignment of the read to the reference. To provide for a gold-standard to compare against, we used BLAST^12^ as a representative of a highly sensitive but slow aligner that likely defines the achievable limits of sensitivity. On synthetic Illumina and PacBio sequencing datasets we noted that GraphMap and BLAST have high precision and recall (∼98%) for both location and alignment measures and are almost indistinguishable in these metrics. Intriguingly, the slight variations in performance that were observed were not defined by the size of the genomes that were studied. In addition, despite the marked difference in error profiles for Illumina and PacBio sequencing, the observed performance metrics were comparable, highlighting the robustness of GraphMap and its similarity to the gold-standard BLAST.

**Figure 2.**
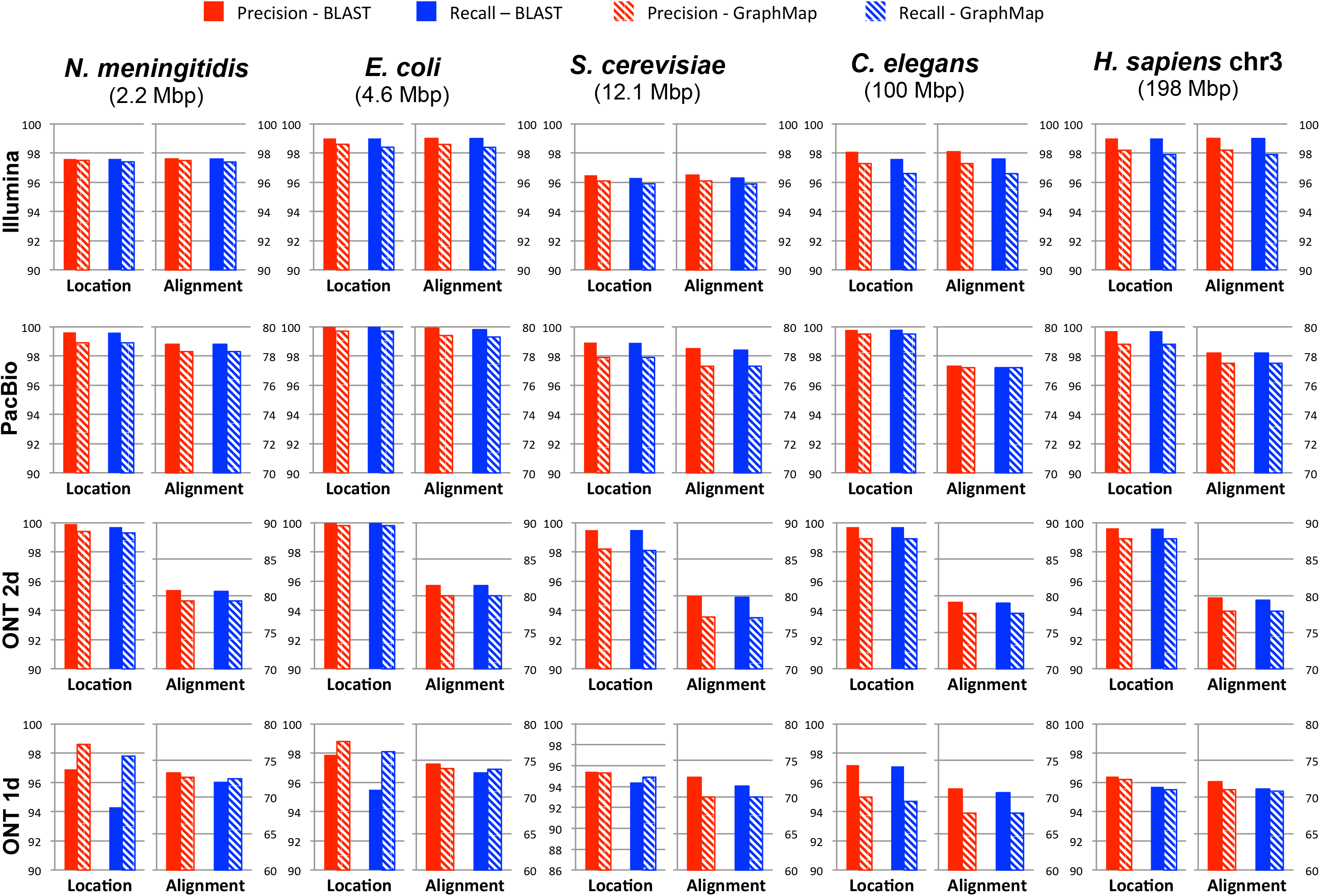
Evaluating GraphMap’s precision and recall against a gold-standard. Results for GraphMap mapping (shaded bars) were determined for a range of genomes (ordered horizontally by genome size from smallest to largest) and sequencing profiles (ordered vertically from low to high error rates) and compared to BLAST (solid bars). For each dataset, the graph on the left shows performance for determining the correct mapping location (within 50 bp; y-axis on the left) and the one on the right shows performance for the correct alignment of bases (y-axis on the right; see **Methods**).

On synthetic ONT data, we noted that slight differences between BLAST and GraphMap were observable but this was less than 3% in the worst case (**Figure 2**). Notably, GraphMap improved over BLAST by similar margins in finding the right mapping location in some cases (e.g. for *N. meningitidis* on ONT 1D data). These slight differences are likely a reflection of the design choices in a sensitive homology detection tool (BLAST) versus a fast and sensitive read mapper (GraphMap), and the impact they have on aligning reads with high error rates. Even with the error rates of ONT 1D data, GraphMap’s precision and recall in selecting the correct mapping location was consistently greater than 95% and 94% respectively. Constructing the correct alignment is more challenging for ONT data as the number of correct bases in the input data is around 70%, but despite this GraphMap correctly aligned ∼70% of the bases. This is likely at the limits of base-level alignment precision and recall as the use of alternate alignment algorithms and parameters did not alter results significantly (**Supplementary Table 1**). Alignments using raw nanopore signal information could be an alternative avenue to boost performance further. These results highlight GraphMap’s ability to identify the precise genomic location based on robust alignments without the need for customizing and tuning alignment parameters to the often unknown error characteristics of the data.

**Table 1.** Comparison of various mappers for single nucleotide variant calling. Results are based on amplicon sequencing data for a human cell line (NA12878) for the genes *CYP2D6*, *HLA-A* and *HLA-B*. Precision values are likely to be an underestimate of what can be expected genome-wide due to the repetitive nature of the regions studied and the incompleteness of the gold-standard set.

|  | **LAST** | **marginAlign** | **BWA-MEM** | **BLASR** | **GraphMap** |
| --- | --- | --- | --- | --- | --- |
| **Precision (%)** | 94 | 100 | 96 | 100 | 96 |
| **True Positives** | 49 | 1 | 47 | 43 | 86 |

While having BLAST-like sensitivity, GraphMap was designed to work with large genomes and sequencing datasets and correspondingly is usually several orders of magnitude faster than BLAST and comparable to other state-of-the-art mappers (**Supplementary Table 1**). For read to reference alignment, BLAST can be feasible for small genomes but can quickly become infeasible for larger genomes (e.g. *C. elegans* or the human genome; **Supplementary Table 1**). GraphMap retains BLAST’s sensitivity while scaling well with genome size. It is possible to tune GraphMap’s settings to make it even faster for short reads with lower error rates, but its sensitivity and speed over a wide range of read characteristics showcases its versatility as a read mapper. Read mappers such as BWA-MEM also exhibit the ability to map varying qualities of reads but need careful tuning of parameters to elicit high sensitivity (**Supplementary Figure 1a**). In addition, for synthetic ONT 1D datasets, we observed a significant drop in precision and recall for read mappers such as BWA-MEM and BLASR in comparison to GraphMap suggesting that they may not be appropriate for such data (**Supplementary Figure 1b, c**). While genome aligners such as LAST can perform better in these settings, they exhibit lower recall for large genomes (a 30% reduction for LAST compared to GraphMap; **Supplementary Figure 1c**) and require significant computational resources for analyzing them (e.g. building the index for a 4 Gbp bacterial genome database with LAST can take more than a week).

**Table 2.** Comparison of various mappers for structural variant calling. Results are based on mapping a MinION dataset for *E. coli* K-12^22^ (R7.3) on a mutated reference containing insertion and deletions in a range of sizes ([100bp, 300bp, 500bp, 1kbp, 1.5kbp, 2kbp, 2.5kbp, 3kbp, 3.5kbp, 4kbp]; 20 events in total). Bold values indicate the best results for each metric. The F_1_ score is given by a weighted average of precision and recall.

|  | **LAST** | **BWA-MEM** | **BLASR** | **GraphMap** |
| --- | --- | --- | --- | --- |
| **Precision (%)** | 0 | 5 | 94 | **100** |
| **Recall (%)** | 0 | 10 | 75 | **100** |
| **$F_1$ Score (%)** | 0 | 6 | 83 | **100** |

### Sensitivity and mapping accuracy on nanopore sequencing data

Encouraged by GraphMap’s performance on synthetic ONT data, we evaluated and compared its results on several published datasets against mappers and aligners that have previously been used (LAST, BWA-MEM and BLASR; see **Methods**). While our synthetic datasets provided a convenient starting point to evaluate performance, they may not capture all features of ONT reads. On the other hand, in the absence of ground truth, evaluating performance using real data can be challenging. To address this, we compared various mappers on their ability to provide accurate (to measure precision of alignments) and complete consensus sequences (as a measure of recall). Overall, the closest competitor to GraphMap was LAST, though it appeared a distant second in terms of these metrics (**Figure 3**). The differences between GraphMap and LAST were apparent even when comparing their results visually, with LAST alignments having low consensus quality even in a high coverage setting (**Figure 3a**). We noted that across datasets, GraphMap mapped the most reads and aligned the most bases, improving sensitivity by 15-80% over LAST and even more compared to other tools (**Figure 3b**; **Supplementary Figure 1**). This led to fewer uncalled bases compared to LAST, BWA-MEM and BLASR, even in an otherwise high-coverage dataset (**Figure 3c, d**). In addition, GraphMap analysis resulted in >10-fold reduction in errors on the lambda phage genome (**Figure 3c**) and reported less than 40 errors on the *E. coli* genome compared to more than a 1000 errors for LAST and BWA-MEM (**Figure 3d**). With ∼80X coverage of the *E. coli* genome, GraphMap mapped ∼90% of the reads and called consensus bases for the whole genome with <1 error in 100,000 bases (Q50 quality). The next best aligner i.e. LAST did not have sufficient coverage (20X) on >7000 bases and reported consensus with a quality of ∼Q36. BWA-MEM aligned less than 60% of the reads and resulted in the calling of >200 deletion errors in the consensus genome. Similar results were replicated in other genomes and datasets as well (**Supplementary Figure 1**). In terms of runtime requirements, GraphMap was typically more frugal than BWA-MEM and slightly slower than LAST (**Supplementary Table 1**).

**Table 3.**
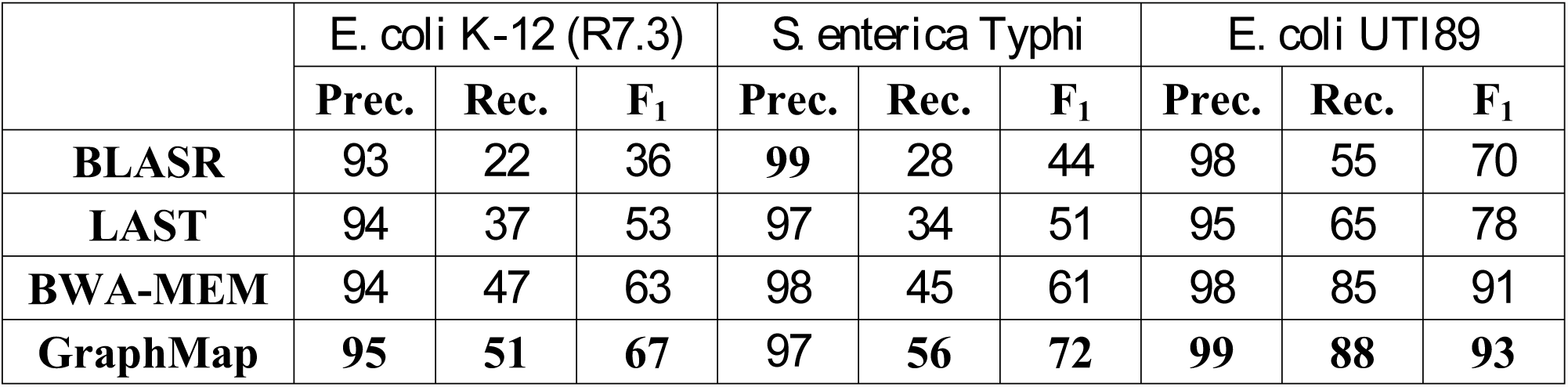
Precision and Recall for species identification using MinION reads. Bold values indicate the best results for each dataset and metric.

|  | *E. coli* K-12 (R7.3) |  |  | *S. enterica* Typhi |  |  | *E. coli* UTI89 |  |  |
| --- | --- | --- | --- | --- | --- | --- | --- | --- | --- |
|  | **Prec.** | **Rec.** | **$F_1$** | **Prec.** | **Rec.** | **$F_1$** | **Prec.** | **Rec.** | **$F_1$** |
| **BLASR** | 93 | 22 | 36 | **99** | 28 | 44 | 98 | 55 | 70 |
| **LAST** | 94 | 37 | 53 | 97 | 34 | 51 | 95 | 65 | 78 |
| **BWA-MEM** | 94 | 47 | 63 | 98 | 45 | 61 | 98 | 85 | 91 |
| **GraphMap** | **95** | **51** | **67** | 97 | **56** | **72** | **99** | **88** | **93** |

**Figure 3.**
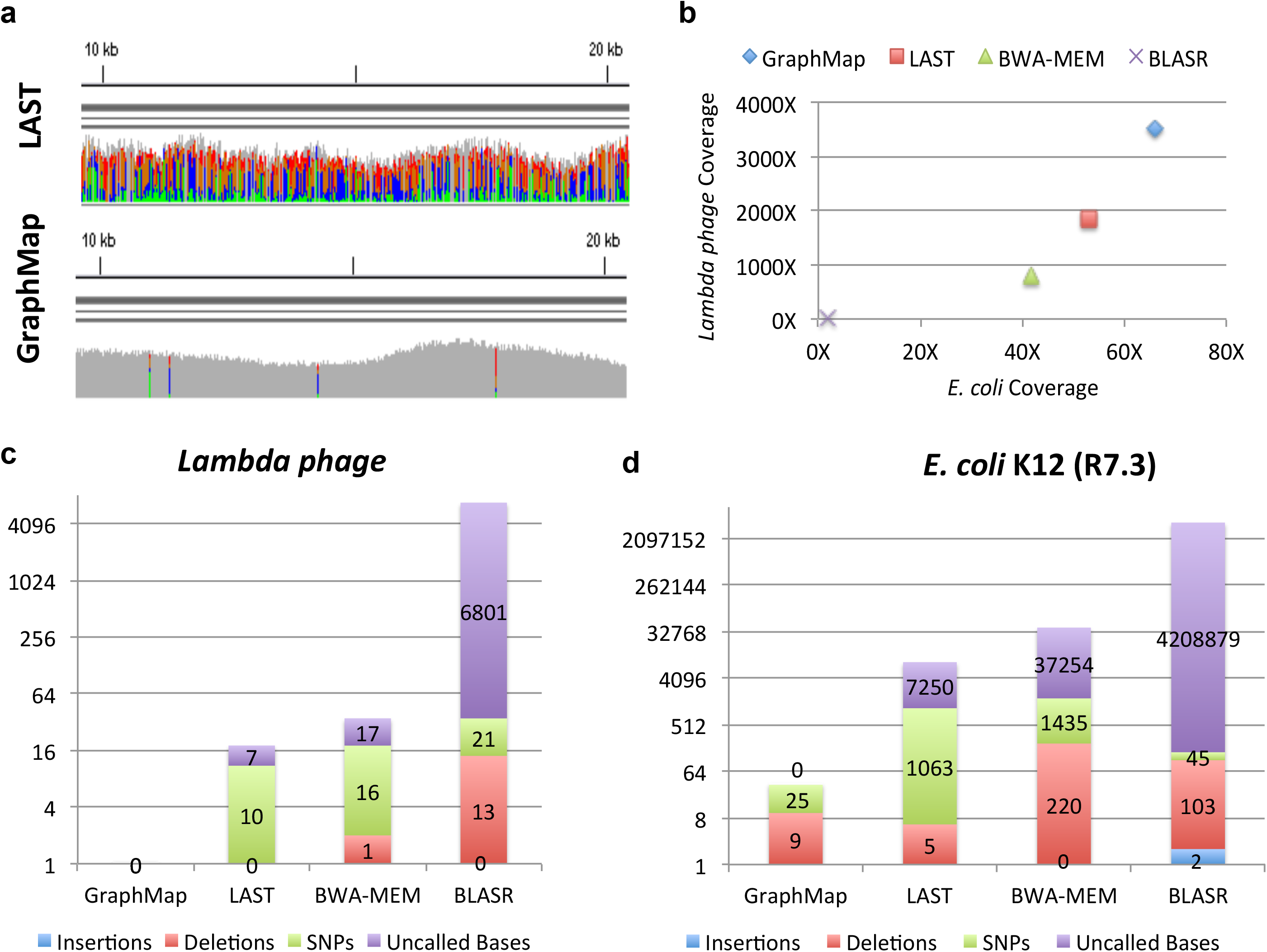
Sensitivity and mapping accuracy on nanopore sequencing data. (a) Visualization of GraphMap and LAST alignments for a lambda phage sequencing dataset on the MinION^8^ (using IGV^26^). Grey columns represent confident consensus calls while colored columns indicate lower quality calls. (b) Mapped coverage of the lambda phage^8^ and the *E. coli* K-12 genome^22^ (R7.3 data) using MinION sequencing data and different mappers. (c) Consensus calling errors and uncalled bases using a MinION lambda phage dataset^8^ and different mappers. (d) Consensus calling errors and uncalled bases using a MinION *E. coli* K-12 dataset (R7.3) and different mappers.

Encouraged by GraphMap’s ability to provide accurate alignments and high quality consensus calls, we used them as a starting point to reanalyze the error profiles of 1D and 2D ONT reads. We reconfirmed substantial variability in the shape and modes of error rate distributions computed by different mappers^2^, but noted that GraphMap’s alignments resulted in lower mismatch rate estimates (**Supplementary Figure 1**). In particular, GraphMap’s distributions were very similar to a maximum-likelihood based realigner (marginAlign^2^), without the need for an expensive realignment step. Overall, deletion and mismatch rates were observed to be higher than insertion rates and significantly reduced from 1D reads (∼15%) to 2D reads (∼7%).

### Application 1: Single-nucleotide variant calling in the human genome with high precision

Variant calling using ONT data has multiple potential hurdles including the lack of a dedicated read mapper or variant caller for it. Not surprisingly, a recent report for calling single nucleotide variants (SNVs) from high-coverage targeted sequencing of the diploid human genome reported that existing variant callers were unable to call any variants and a naive approach requiring 1/3 of the reads to support an allele could lead to many false positive variants^13^. To evaluate if improved read mappings from GraphMap could increase sensitivity and precision, we reanalyzed data reported in Ammar et al^13^, adopting a variant caller (LoFreq) that directly utilizes information about base qualities for being robust to high error rates^3^. We used a set of confident calls for this sample (NA12878) as our gold standard^14^. Our results provide the first demonstration that nanopore data can be used to call heterozygous variants in challenging regions of the human genome (in the genes *CYP2D6*, *HLA-A* and *HLA-B*) with high precision (>96% with GraphMap; **Table 1**). These can then be the foundation for reconstructing haplotypes, in these complex but clinically important regions of the human genome, by exploiting the advantages of long spanning reads. Significantly, we noted that alignments from GraphMap provided many more true positives than the next best mappers (BWA-MEM, LAST) providing a 76% improvement in recall overall (**Table 1**). Confirming the report in Ammar et al^13^, our results suggest that targeted nanopore sequencing reads can be mapped to the correct location on the human genome despite the presence of very similar decoy locations (94% identity between *CYP2D6* and *CYP2D7*), with GraphMap providing the most on-target reads (**Supplementary Figure 1**).

**Figure 4.**
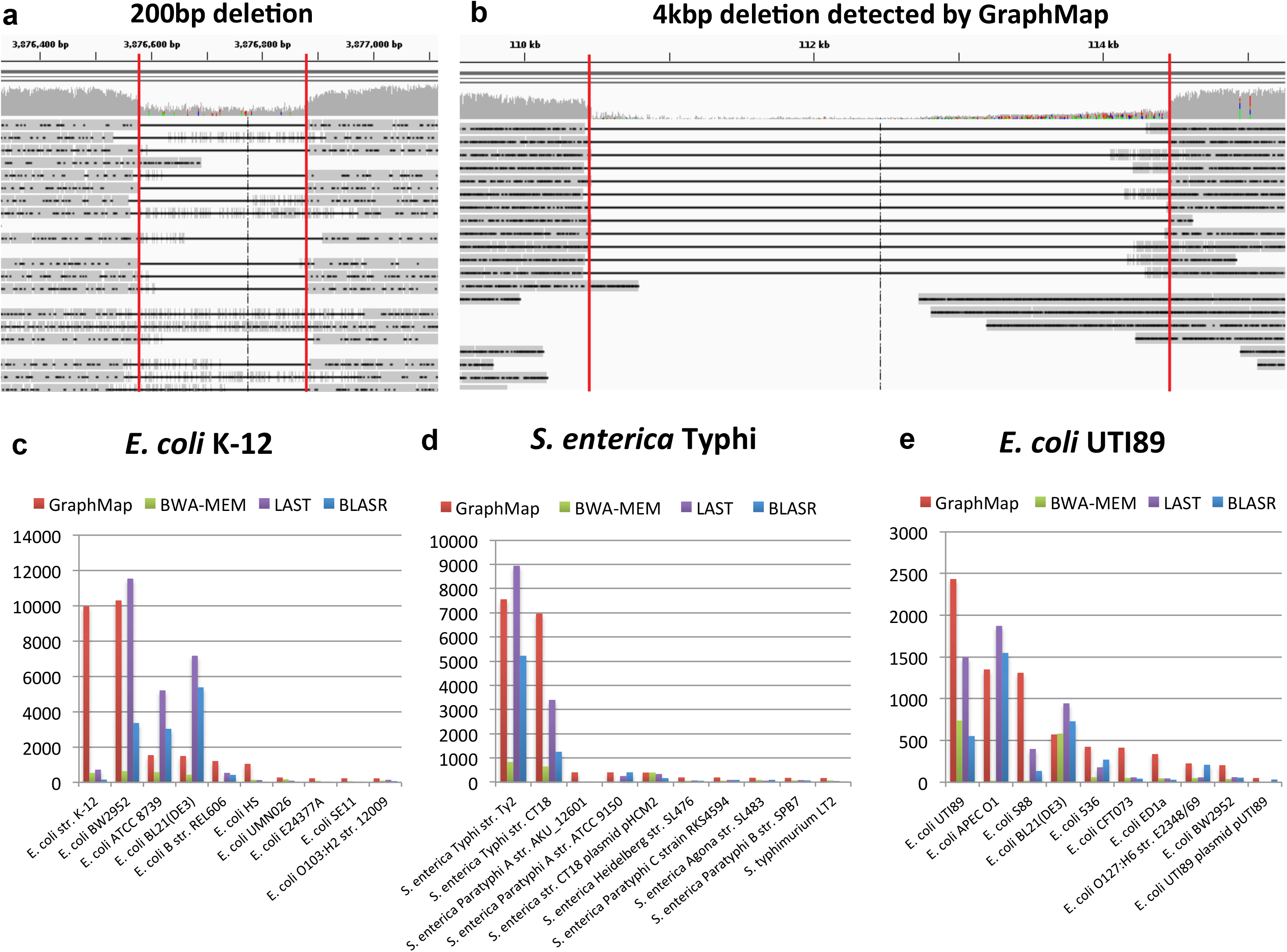
Variant calling and species identification using nanopore sequencing data and GraphMap. (a) An IGV view of GraphMap alignments that enable the detection of a 200bp deletion (delineated by red lines). (b) GraphMap alignments spanning a ∼4 kbp deletion (delineated by red lines). Number of reads mapping various genomes in a database (sorted by GraphMap counts and showing top 10 genomes) using different mappers (GraphMap, BWA-MEM, LAST and BLASR) and three MinION sequencing datasets for (c) *E. coli* K-12^22^ (R7.3) (d) *S. eneterica* Typhi and (e) *E. coli* UTI89. Note that GraphMap typically maps the most reads to the right reference genome (at the strain level) and the *S. eneterica* Typhi dataset is a mixture of sequencing data for two different strains for which we do not have reference genomes in the database.

### Application 2: GraphMap enables sensitive and accurate structural variant calling

Long reads from the MinION sequencer are, in principle, ideal for the identification of large structural variants (SVs) in the genome^15^, but this has not been explored before with the limitations of existing tools^1^. Using real *E. coli* data mapped to a mutated reference we systematically evaluated this application and observed that GraphMap’s alignments could readily detect SVs, both insertions and deletions, over a range of event sizes (100bp-4kbp; **Table 2**). Furthermore, GraphMap produced alignments that accurately demarcated the alignment event and did this without reporting any false positives (**Figure 4a,b** and **Table 2**). These alignments provided perfect recall over the entire range of indel sizes (100bp-4kbp) and a 35% improvement in recall over the next best mapper (which was BLASR for this application). Highlighting the non-trivial nature of this problem, BWA-MEM alignments resulted in low precision and recall (≤10%), with many false positives, while LAST alignments were unable to detect any of the events under a range of parameter settings (**Table 2**). These results emphasize GraphMap’s advantage for the purpose of systematically cataloging point mutations as well as structural variations using nanopore sequencing data.

### Application 3: Sensitive and specific pathogen identification with ONT reads

Due to its form factor and real time nature, an application of MinION sequencing that has garnered interest in the community is in the identification of pathogens in clinical samples. Sequencing errors (particularly in 1D data) and the choice of read mapper could significantly influence results in such an application and lead to misdiagnosis. GraphMap’s high specificity in read mapping as seen in the results for Ammar et al (**Supplementary Figure 1**) suggested that it could be useful in this setting. Using clonal sequencing data on the MinION and a database of microbial genomes we created several synthetic benchmarks to evaluate the performance of various mappers for this application (see **Methods**). For species level identification, we noted that all mappers reported high precision (typically >95%) but recall varied over a wide range from 20% to 90% (**Table 3**). GraphMap had the highest recall and F_1_ score in all datasets providing an improvement of up to 10% over other mappers. For this application, BWA-MEM was the next best mapper while LAST and BLASR exhibited 20% reduced recall compared to GraphMap (**Table 3**). Not surprisingly, strain level identification using MinION data appears to be much more difficult and in some cases a closely related strain can attract more reads than the correct strain (**Figure 4c**). However, in the datasets that we tested we noted that GraphMap assigned most reads to a handful of strains that were very similar to the correct strain (**Figure 4c- e**; 99.99% identity for *E. coli* K-12 and BW2952). Moreover, the use of strain specific sequences was able to unambiguously identify the correct strain from this subset (e.g. there were no reads mapping to NC_012759.1:4.13Mbp-4.17Mbp, a region unique to BW2952), suggesting that this approach could be used to systematically identify pathogens at the strain level.

## Discussion

The development of GraphMap provides a new opportunity in the tradeoff between mapping speed and sensitivity. It demonstrates BLAST-like sensitivity while being comparable in speed to other state-of-the-art short and long-read mappers. On recently available nanopore sequencing data, GraphMap is unmatched in terms of sensitivity, mapping more than 90% of reads and bases on average. Our comparisons with BLAST suggest that reads that cannot be mapped by GraphMap may essentially be unmappable. High sensitivity is a key requirement for mapping tools as typically reads that cannot be mapped are lost from downstream analysis.

GraphMap’s speed and sensitivity do not come at the expense of location and alignment precision, as demonstrated by our experiments with synthetic and real datasets. For determining the correct genomic location, GraphMap’s precision is typically greater than 98% and it is able to distinguish between candidate locations that are more than 94% identical on the human genome. For alignment precision, GraphMap’s performance scales according to sequencing error rate, is comparable to BLAST and was observed to be robust to choice of alignment algorithms and parameters. It should thus provide a better starting point for downstream analysis tools including realigners and consensus calling algorithms such as marginAlign^2^ and Nanopolish (https://github.com/jts/nanopolish).

Applications such as variant calling and species identification can be challenging with PacBio and nanopore sequencing data, due to ambiguities in mapping and alignment. We show that despite the lack of custom variant callers, read mappings from GraphMap can lead to sensitive and precise variant calls. Particularly exciting is the ability to call structural variations over a range of event sizes without having to assemble the reads. No doubt, the development of new nanopore-specific tools is likely to improve the quality and precision of structural variant calls even further, but GraphMap alignments can provide a useful starting point for such applications. We also observed that GraphMap alignments could be used to identify the species-level origin of reads with high precision and recall. The sensitivity of mapping with GraphMap can be a key advantage in applications where MinION sequencing reads are used in real-time to identify pathogens^16^. With further downstream processing, these read mappings could be used for strain-level typing and characterization of antibiotic resistance profiles^16^, meeting a critical clinical need.

In principle, the approach used in GraphMap could be adapted for the problem of computing overlaps and alignments between reads. As was recently shown, nanopore sequencing reads can be used to construct high-quality assemblies *de novo*^17^. GraphMap’s sensitivity and specificity as a mapper could thus serve as the basis for fast computation of overlap alignments and *de novo* assemblies in the future.

## Methods

### Description of the GraphMap Algorithm

#### Stage I: Region selection

GraphMap starts by roughly determining regions on the reference genome where a read could potentially be aligned. This step is performed in order to reduce the search space for the next step of the algorithm, while still providing very high sensitivity. As a first step, region selection relies on finding seeds between the query sequence and the reference, before clustering them into candidate regions. For seed finding, we found that commonly used approaches such as maximal exact matches (MEMs) (as used in BWA-MEM^6^) or Hamming distance based spaced seeds (as used in LAST^10^) are either not sensitive enough or not specific enough in the presence of error rates as high as is feasible in nanopore data. Instead, we employed a form of gapped spaced seeds similar to gapped q-gram filters for Levenshtein distance^11^. Specifically, we extended the approach proposed in Burkhardt and Kärkkäinen^11^ to use both one- and two-gapped q-grams (**Figure 1b**) as detailed below. This allows us to accommodate an arbitrary number of gaps in the q-gram.

Gapped q-grams are a seeding strategy that allow for fast and very sensitive lookup of inexact matches, with variations allowed in predefined “don’t care” (DC) positions of the seed. Concordant with existing terminology, we call the concrete layout of the inclusive and DC bases a shape and the number of used positions its weight. Gapped q-grams allow for DC positions within a shape to also contain insertions and deletions (indels). Our approach for implementing Levenshtein gapped q-grams is based on constructing a hash index of the reference sequence, where the q-gram positions are hashed by the keys constructed from the shape’s layout – only inclusive bases are taken for constructing the key, while the DC bases are simply skipped (**Figure 1b)**. During the lookup step, multiple keys are constructed for each shape and used for retrieval. For each DC base, three lookup keys are constructed:

I. A key constructed in the same manner as during the indexing process, which captures all seeds with a DC base being a match or a mismatch (e.g. “1110111”),
II. A key where the DC base is not skipped. This key captures up to one deletion at the specified position (e.g. “111111”), and
III. A key where the DC base as well as the following base is skipped. This key allows for at most one insertion and one match/mismatch (e.g. “11100111”).

In total, for each shape *d*^3 keys are constructed, where d is the number of DC bases. GraphMap uses two shapes for the region selection process: “1111110111111” (or the 6-1-6 shape) and “11110111101111” (or the 4-1-4-1-4 shape), where 1 marks the inclusive bases and 0 the DC positions. This shape combination was selected based on empirical evaluation of a range of combinations, due to the computational intractability of computing the optimal shape for the Levenshtein distance. For each shape, a separate index is used. At every seed position, both shapes are looked up, and all hits are used in the next step for binning.

To derive a general approach for binning seed hits, we draw on the concept of a Hough Transform (HT), a method commonly used in image processing for detection of shapes such as lines, circles and ellipses. The HT defines a mapping from image points into an accumulator space, called the Hough space. In the case of line detection, if a given set of points in Cartesian space are collinear, then their relation can be expressed with a line equation with common slope *m* and intercept *c*:

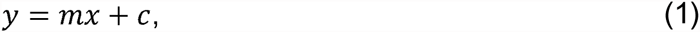

where (*x*, *y*) are the coordinates of a point in 2D space. HT attempts to determine parameters m and c of a line that describes the given set of points. One must note that the system is generally overdetermined (whenever there are more than two points given), and thus the problem can be solved using linear regression techniques. However, the HT uses an evidence-gathering approach, which can be used to detect an arbitrary number of lines in the image instead of only one best. Equation (1) can be converted into its dual in parameter space:

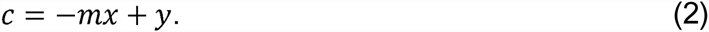

The intuition is as follows: given a point (*x*, *y*) in Cartesian space, its parameter space representation defines a line. If multiple Cartesian space points are given, each transforms into a different line in the parameter space. Their intersections specify potential lines in the original, Cartesian space. HT defines an accumulator space, in which m and c are rasterized so as to take only a finite range of values. HT then simply counts all the potential solutions in the accumulator space by tracing all the dual lines for each point in the Cartesian space, and increasing the vote count for each (m, c) coordinate. All HT space coordinates with count above a defined threshold can then be considered as candidate lines in the original Cartesian space.

A single seed hit can be represented with a *k-point* (*q*, *t*) in 2D space, where q is the seed’s position on the read, and t is the position of the seed hit on the reference. In the case a read is completely error-free and extracted from the exact reference, its set of k-points would be perfectly collinear in such defined space. Moreover, under these ideal conditions, they would all lie on a line tilted at a 45° angle (slope *m* = 1). This collinearity also corresponds to the main diagonal in the dynamic programming alignment matrix. Since *m* is known, only the intercept parameter c needs to be determined to find the accurate mapping position. For this, the HT voting mechanism can be used. Again, since *m* is known, the 2D accumulator space is not required – only an array for the c value is sufficient. As *c* corresponds to the (already discrete) coordinates on the reference sequence, a simple integer array of the length of the reference can be used for counting votes. For each k-point, its *c* parameter value is determined with a simple expression:

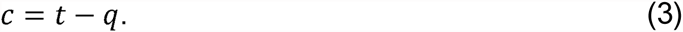

The index of the accumulator array with the highest count is the exact mapping position of the read on the reference. In this simple form, this approach mirrors the techniques used in other aligners (e.g. FASTA). However, the concept of the Hough Transform (HT) allows us to extend and generalize this notion.

We account for substitution and indel errors in this framework as follows: substitution errors cause only the reduction in the maximum vote count for the correct c value and induce noise votes in other locations on the reference. Such type of errors can be addressed using appropriate thresholding on the hit count (see below). On the other hand, indels are of special interest because they shift the alignment diagonal and cause more substantial reduction of votes for the correct location. Additionally, using an accumulator array that is of size equal to the size of the reference sequence can cause high memory consumption, especially in the case of processing large sequences in multithreaded environments.

To address both the error-rate and memory consumption issues, we rasterize the reference sequence into partitions of length *L/3*, where *L* is the read length. For each seed hit, we increase the value of the bin corresponding to its c parameter value determined using Equation (3). If a bin has multiple hits from the same seed, only one hit is counted. Bins are then sorted in descending order of the number of hits. Only bins which have a count ≥0.75. *b*_*max*_ are selected for further processing, where *bmax* is the count of the highest scoring bin. We then define a *region* as a portion of the reference that expands the corresponding bin’s start and end location by an additional read length, to compensate for potential indel errors and ensure that the entire alignment area enters the next step of mapping. In case the reference genome has been specified as being circular by the user and a selected region should span beyond any end of the reference, then the region is constructed by concatenating the beginning and the end of the reference sequence. Regions are then processed separately until the last step of the method, when the highest scoring region is selected for alignment.

#### Stage II: Graph-based vertex-centric construction of anchors

In this stage, we attempt to refine candidate regions from stage I by constructing alignment chains or *anchors* from short seeds matches. To do this, we introduce the notion of a *kmer mapping graph*. Given a pair of sequences (target and query), the method starts by constructing a kmer mapping graph from the target sequence. Target and query sequences in this case are the read sequence and a single region of the reference sequence. Whether the read or the region is chosen to be the target sequence is not essential for the approach to work, but in our implementation, we chose the read to be the target sequence in order to reduce memory consumption (the region determined is always larger than the read). The vertices of the kmer mapping graph are the kmers of the target sequence of length *T*. Unlike the *de Bruijn* graph, identical kmers are not truncated into the same vertex of the graph but are kept as separate individual vertices. For every vertex *vi* (∀i∈ (0…*T* - k)), *l* directed outbound edges are added which connect *v*_*i*_ to vertices *v*_*i*+1_, *v*_*i*+2_,…, *v*_*i*+l_. Note that, since directed edges are added in this consecutive manner, the last (*l* - 1) vertices of the graph cannot have *l* outbound edges since there are no vertices to connect them to in the graph. In summary, a kmer mapping graph is a directed acyclic graph, consisting of (*T* - *k* + 1) vertices and 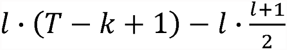 directed edges.

The rationale for such a design is as follows. In case *l* = 1 and if the query is a subset of the target with no differences or errors, the target’s mapping graph would contain the same kmers in the exact same order as can be found in the query sequence the read originated from. Thus, an exact walk exists in both sequences. However, in realistic conditions, variations and sequencing errors exist in reads. Although the majority of kmers might still be in the same order, a simple exact linear walk through the reference’s and read’s mapping graphs cannot be found due to the differing kmers present. Instead, the walk is fragmented into several smaller ones. The fragmentation is especially large in data with high error rates, such as those obtained with nanopore sequencing. In these cases, it is sometimes difficult to find even two consecutive correct kmers. To address this issue, the additional (*l* – 1) edges act as a *bridge* between vertices in the mapping graph. Thus, we allow a linear walk to be found not only by following consecutive kmers in the graph, but to jump-over those that produce poorer solutions. **Figure 1d** depicts such an example.

In order to find an appropriate mapping position, we conduct a simultaneous walk both in the target sequence and in the query. The mapping graph is constructed from only the target sequence, while the walk in the query is conducted by iterating through all its consecutive kmers. In the mapping graph, all walks that correspond to potential mapping sites are simultaneously monitored and extended. Although this approach may appear to be time-consuming, GraphMap handles it in an elegant and efficient vertex-centric manner as detailed below.

Note that at this stage of the algorithm, GraphMap does not use the same index as in the region selection process. Instead, a new index is constructed from the target on the fly, using a much smaller seed size (default is *k* = 6). In principle, an arbitrary indexing method such as suffix arrays, FM index or hashing can be used at this stage. In our implementation, perfect kmer hashing is used for indexing when *k* < 10 and otherwise suffix arrays are used. Following graph construction, the next step is to do graph traversal. For each consecutive kmer in the query, a list of hits on the target sequence is obtained from the index. For every hit, its position on the target directly points to the vertex in the graph that the kmer belongs to. The vertex-centric walk can then elegantly be described as follows: for a chosen vertex, collect information from its input edges, choose the “best” edge and update the information it contains, and transmit this information to all outbound edges simultaneously. We define the “best” edge to be the one belonging to the longest walk. The information that is transmitted through the edges contains the walk length, the position of the starting kmer in both the target and the read, and the number of covered bases and kmers in both sequences. Consistently transmitting this information only requires updating the current walk length, number of kmers and the number of covered bases. Since the only operations we perform on a vertex include simple collect, extend and transmit steps, the runtime complexity of the vertex-update operation is O(1). Initially, the information stored in the graph is set so that there are no valid walks (all walk lengths are set to zero). The walk through the graph can then be viewed as propagating the information through the directed acyclic graph from the initial candidate location of the mapping to the furthest reachable vertex.

After all kmers from the query have been processed, a list of walks in the graph is collected. Walks which contain less than a user defined amount of covered bases in both sequences (default of 12 bases) are not processed further. Each walk represents one valid candidate location for mapping of the query. Intuitively, one would simply choose the longest walk for the most probable mapping position. Our empirical tests suggest that it is possible to achieve relatively high accuracy even with this simple heuristic. Indeed, in the presence of low substitution error rates (as is the case for Illumina as well as PacBio reads), a single walk can cover most of, if not the entire read. However, accuracy can greatly be impaired in the presence of higher substitution error rates as seen in nanopore sequencing data. In this case, clusters of errors in reads tend to cause fragmentation of walks in the mapping graph, resulting in a list of shorter walks (typically several kmers long), none of which span more than a small percentage of the read length. Although these small walk fragments do not seem to carry much information, they actually represent an exact ordering of kmers in both sequences and thus form the basis of a longer alignment. We refer to these short walks as anchors for simplicity, although they differ from the traditional definition of an anchor in that walks allow for mismatches and indels to be present within them.

#### Stage III: Extending anchors into alignments using LCS

Each anchor reported by GraphMap in stage II represents a shared segment (or subsequence) between the target and the query sequence with known start and end positions in both sequences. Due to the presence of repeats, the set of anchors obtained is not necessarily monotonically increasing in both the target and query coordinates. For this reason, a subset of anchors that satisfy the monotonicity condition needs to be selected. The problem of identifying such a subset can be expressed as finding the Longest Common Subsequence in k Length Substrings^18^ (LCSk). Recently, an efficient and simple algorithm for solving a variant of the LCSk problem has been proposed^19^. In our implementation we follow this paradigm and instead of using substrings of fixed size k, we allow for variable length substrings. Concretely, the size of each substring is equal to the length of the corresponding anchor in both sequences. As a result, the reconstruction of LCSk is obtained in the form of a list of consecutive anchors in the target and the query sequence.

#### Stage IV: Refinining alignments using L_1_ linear regression

The alignments obtained using LCSk tend to be largely accurate but since its definition lacks constraints on the distance between substrings, the alignments obtained may include outlier matches and mis-estimation of overall alignment length (**Figure 1e)**. These outliers are caused by repeats or sequencing errors, but they still satisfy the monotony condition. Similar to the observation we presented in the region selection step, the LCSk list of anchors should ideally be collinear in the 2D query-target coordinate space, with a slope of 45°. All deviations from this line are caused by indel errors, and can be viewed as noise. We start the filtering of the LCSk outlier anchors by fitting a 2D line with a 45° slope in the query-target space under the least absolute deviation criteria (LAD, L_1_). Next, a subset of anchors which are located within 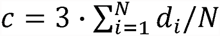 from either side of the L_1_ line is selected, where e is the expected error rate (by default, conservatively set to 45%), r is the target (read) length, and the factor 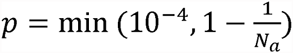 is used to convert the distance from target coordinate space to a distance perpendicular to the L_1_ line. A confidence interval 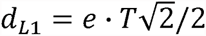 is calculated, where di is the distance from a selected anchor i to the L_1_ line. LCSk is then repeated once again but only on the anchors which are located within the distance ±c from the L_1_ line in order to compensate for possible gaps caused by anchor filtering.

After filtering, five scores that describe the quality of the region are calculated. They include: the number of exact kmers covered by the anchors *nkmers*, the standard deviation σ of anchors around the L_1_ line, the length of the query sequence which matched the target (distance from the first to the last anchor) mlen, the number of bases covered by anchors (includes only exact matching bases) ncb and the read length. The last four scores are normalized to the range [0,1] with the following equations (4)-(7):

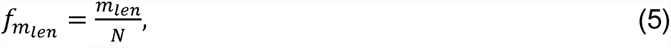

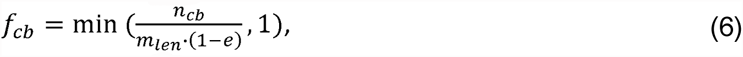

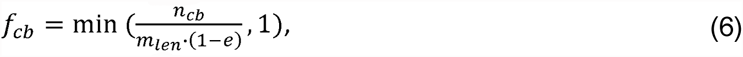

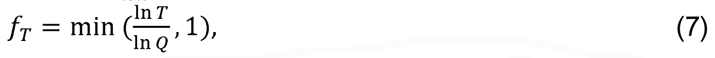

where *Q* is the length of the reference sequence (query in our previous definition). The overall quality of the alignment in a region is then calculated as:

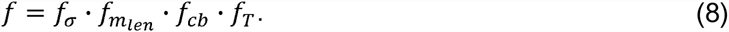

#### Stage V: Construction of final alignment

After all selected regions have been processed, they are sorted by the *f* parameter. The region with the highest value *fmax* is selected for the final alignment. Unlike many other methods which use the seed-and-extend approach, GraphMap aligns the entire read using the semi-global alignment algorithm. The default parameters of GraphMap use an implementation of Myers’ bit-vector approach for alignment^20^ with alignment parameters fixed to match = 1, mismatch = -1, gapopen =0 and gapextend = -1. GraphMap also allows users a choice of aligners and custom scoring parameters. Current alternative alignment options include an implementation of Gotoh’s semi-global alignment algorithm^21^ as well as an option to construct anchored alignments. Specifically, in the anchored approach, anchors from the LCSk step are clustered and alignments within and between cluster endpoints computed using Myers’ bit-vector alignment (extensions to read ends are done without gap penalty). Clustering is done by collecting neighboring anchors where the ratio of distances in the read and reference coordinates is less than *e*/2 (as before, *e* is the expected error rate in the data). Clusters with very few bases (<30 or 2% of read length) were discarded for this purpose.

GraphMap allows users to output all equally or similarly good secondary alignments by specifying an ambiguity factor *F* in the range [0,1] and using that to select regions which have *n*_*kmers*_ ≥ (1- *F*) · *n*_*kmers,best*_, where *n*_*kmers,best*_ is the number of kmers of the region with the maximum f value. We denote the count of regions with *n*_*kmers*_ above the ambiguity threshold as *N*_*a*_.

#### Mapping quality

Since the region filtering process in GraphMap maintains a large collection of possible mapping positions on the given reference, it enables meaningful calculation of the mapping quality directly from its definition:

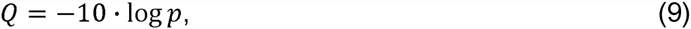

where *p* is the probability of the read being mapped to the wrong position. We calculate *p* simply as 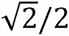 and report quality values according to the SAM format specification.

#### E-value

For each reported alignment, GraphMap calculates the *E*-value which is given as a custom “ZE” parameter in the output SAM file. Following the approach used in BLAST, we rescore alignments and use pre-calculated Gumbel parameters to compute *E*-values in the same way as in BLAST (default scoring parameters: *match* = 5, *mismatch* = -4, *gap*_*open*_ = -8 and *gap*_*extend*_ = -6).

### Datasets

For evaluating GraphMap and other tools, we used five publicly available MinION sequencing datasets, 20 synthetic datasets and MinION sequencing reads for an *E. coli* UTI89 sample as detailed below.

#### MinION library preparation

Genomic DNA was extracted from *Escherichia coli* UTI89 using the QIAamp® DNA mini kit (Qiagen). 1μg of the extracted DNA was then sheared in a total volume of 80μl using a Covaris g-TUBE according to the manufacturer’s instructions with centrifugation for 1min at 6000rpm. Sheared DNA was end repaired and A-tailed using the GeneRead™ DNA Library Prep I Kit from Qiagen according to the manufacturer’s protocol. The reaction was purified using 1X volume of Agencourt Ampure XP beads and eluted in 30μl nuclease-free water. Subsequent steps of DNA sequencing library preparation were carried out using Oxford Nanopore’s MinION Genomic DNA Sequencing Kit (SQK-MAP003) according to the manufacturer’s recommended protocol, including the addition of purified BSA (NEB) to Agencourt Ampure XP beads and Elution buffer.

#### MinION sequencing of E. coli UTI89

Immediately prior to sequencing, 12μl of the DNA library was combined with 134μl EP buffer and 4μl Fuel Mix and mixed by inversion 10 times. The flow cell was primed by washing with two aliquots of 150μl of EP buffer, with ten minutes in between washes. 150μl of the prepared DNA Library was then loaded onto the flow cell and the Genomic DNA 48 hour sequencing run program was selected. Fresh sample was loaded onto the flow cell at 12 hour intervals throughout the run.

#### Publicly available sequencing datasets

Five publicly available MinION sequencing datasets were used for evaluation. These included a lambda phage dataset, two *E. coli* datasets (each produced with a different version of MinION chemistry), a *S. enterica* Typhi dataset and a dataset consisting of three amplicons from the human genome:

**(I) Lambda phage burn-in dataset^8^**. The dataset consists of 40,552 reads in total (211 Mbp of data), generated using an early R6 chemistry. The reference genome (NC_001416) is 49 kbp long giving an expected coverage of >4300X.

**(II) *E. coli* K-12 MG1655 R7 dataset^22^**. The dataset has 111,128 reads (668 Mbp) providing 144X coverage of a 4.6 Mbp genome (U00096.2).

**(III) *E. coli* K-12 MG1655 R7.3 dataset^22^**. The dataset has 70,531 reads (311 Mbp) providing 67X coverage of the genome (U00096.2).

**(IV) *S. enterica* Typhi dataset^1^**. The dataset is composed of two runs of strain H125160566 (16,401 reads and 6,178 reads respectively) and one run of strain 08-04776 (10,235 reads).

**(V) Amplicon sequencing of human *HLA-A*, *HLA-B* and *CYP2D6* genes^13^**. The dataset contains 36,779 reads in total.

#### Synthetic datasets

Synthetic Illumina reads were generated using the ART simulator^23^ (150 bp single-end) and PacBio CLR reads using the PBSIM simulator^24^ (with default settings). For synthetic MinION data we adopted PBSIM (as no custom ONT simulators exist currently) and used parameters learnt from LAST alignments (to avoid bias towards GraphMap) with *E. coli* K-12 R7.3 data (**Supplementary Table 4**). Reads were simulated for five reference sequences: *N. meningitidis* serogroup A strain Z2491 (1 chromosome, 2.2 Mbp, NC_003116.1), *E. coli* K-12 MG1655 (1 chromosome, 4.6 Mbp, U00096.2), *S. cerevisiae* S288C (16 chromosomes, 12 Mbp), *C. elegans* (6 chromosomes, 100 Mbp) and *H. sapiens* Chr3 (198 Mbp, hg19 v38, CM000665.2).

### Evaluation methods

#### Performance on synthetic data

Mappers were evaluated for precision and recall in meeting two goals:

1. Finding the correct mapping location – a read was considered correctly mapped if its mapping position was within 50bp of the correct location.
2. Reporting the correct alignment at a per-base-pair level – a base was considered correctly aligned if it was placed in exactly the same position as it was simulated from.

#### Parameter settings for mappers

We evaluated BWA-MEM using the nanopore setting (-x ont2d; version: 0.7.10-r1027-dirty) and for detecting structural variations we increased the alignment bandwidth using “-w 5000 -d 5000”. BLASR was evaluated with the options “-sam -bestn 1” (version: 1.3.1) and in addition for the database search we set more stringent parameters (“-minMatch 7 -nCandidates 1”). LAST was run with a commonly used set of nanopore settings^22^ (“-q 1 -r 1 -a 1 -b 1”) and with additional overlap mode setting (“-T 1”; to force end-to-end alignment) for structural variant detection (version: 475). BLAST (version: ncbi-blast-2.2.30+-x64-linux) was run with default settings for Illumina data and a more suitable nanopore setting^25^ “-reward 5 -penalty -4 -gapopen 8 -gapextend 6 -dust no” for ONT and PacBio data. GraphMap (version: v0.21) was run with default settings. In addition, for circular genomes we used the -C option, anchored alignments for calling structural variations (“-a anchor”) and *E*-value filtering (“-z 1e0”) for database search and variant calling. We used marginAlign with the parameter “--em” for variant calling (version: (Git commit) dfdb05d6d291aab186b6f3668fa3d7c1de28787d).

#### Consensus calling using MinION data

Consensus was called using a simple majority vote of aligned bases, insertion and deletion events (insertion sequences were taken into account while counting events) and positions with <20X coverage were not called. All reads were mapped and analyzed to determine consensus sequences.

#### Benchmarking mappers for pathogen identification

Bacterial genomes similar to a list of water-borne pathogens were selected from NCBI’s bacterial database to construct a database of 268 genomes (550 Mbp; **Supplementary Table 5**). MinION sequencing datasets from cultured isolates were used as proxy for sequencing of pathogen-enriched clinical samples (using data for *E. coli* K-12 R7.3, *S. enterica* Typhi and *E. coli* UTI89, as specified earlier). This is a simple test case as real samples are likely to have contaminations from other sources as well (e.g. human DNA). We mapped these three read datasets to the database of bacterial genomes using each of the mappers to find unique alignments and test if these could help identify the correct species and strain. For BWA-MEM and LAST, we chose the best alignment based on alignment score (as long as alignment score and mapping quality were greater than 0) and for GraphMap and BLASR we used the unique reported alignment (mapping quality > 0).

#### Single nucleotide variant calling

All 2D reads from Ammar et al^13^ were mapped to the human genome (GRCh37.p13; chr 6 and 22) and for each read only the alignment with the highest alignment score (AS) was kept. To avoid chimeric reads as reported in the original study we used only reads that fully spanned the amplicon regions for this analysis. Variants were called using LoFreq^3^ with the parameters “-a 0.01 -q 0 -Q 0 --no-default-filter”. We then compared the detected SNVs with known variants from dbSNP and a high-confidence set for NA12878^14^ (the HapMap sample used for sequencing; ftp://ftp.ncbi.nlm.nih.gov/snp/organisms/human_9606_b141_GRCh37p13/VCF/All.vcf.gz; ftp-trace.ncbi.nih.gov/giab/ftp/data/NA12878/variant_calls/NIST/NISTIntegratedCalls_14datasets_131103_allcall_UGHapMerge_HetHomVarPASS_VQSRv2.18_all.primitives.vcf.gz) to identify true positives and false positives.

#### Structural variation detection

We modified the *E. coli* K-12 MG1655 reference by inducing 20 SV events (10 insertions and 10 deletions) of different sizes: 100bp, 300bp, 500bp, 1000bp, 1500bp, 2000bp, 2500bp, 3000bp, 3500bp, 4000bp. All 2D reads from both *E. coli* K-12 datasets (R7 and R7.3) were combined and mapped. SVs were detected by simple consensus vote of indel events reported in the alignments (≥20 bases to avoid sequencing errors). Note that a realigner such as marginAlign is not designed for this application and hence we did not evaluate its use here. In the absence of a sophisticated SV caller for nanopore data we used a simple rule that identifies windows where >15% of the reads at each position report an insertion (or deletion) event (at least 5 reads). To avoid fragmented events due to a local drop in allele frequency, we merged windows which were less than window-length apart (max of the two windows). We considered a detected event a true positive if its size was within a 25% margin of the true size and its start and end locations were less than 25% of event size away from the true locations.

